# Impaired Associative Memory, Inference, and Theta Dynamics in Postictal Psychosis of Epilepsy

**DOI:** 10.64898/2026.01.23.701370

**Authors:** Aryeh Dworkin, Danying Wang, Diego Jiménez, Charlotte Ravenscroft, Francesco Turco, Chloe Johnson, Fahmida Chowdhury, Joao Pizarro, Matthew Walker, Simona Balestrini, Daniel Bush, Umesh Vivekananda

## Abstract

Postictal psychosis (PIP) is a severe complication occurring in 2% of people with epilepsy (PWE) whose underlying pathophysiology remains poorly understood. Although historically considered separate from other forms of psychosis, newer evidence demonstrates a shared genetic susceptibility. People with schizophrenia are typically impaired at both associative learning and inferring connections between overlapping associations. Successful associative encoding, retrieval, and inference can each be predicted by changes in frontotemporal theta band activity, which is impaired in rodent models and people with schizophrenia. Here, we recorded high-density scalp EEG from PWE with history of PIP and well-matched control participants while they undertook a memory inference task. We found that associative memory and inference were both impaired in the PIP group, despite no difference in item recognition. Moreover, we found disrupted theta activity during memory encoding and the retrieval of inferred associations in PWE with PIP that likely originated from the medial temporal and frontal lobes. These results suggest a pattern of behavioural deficits and altered neural dynamics common to both PIP and schizophrenia. Interpreted in conjunction with previous genetic studies, they may reflect shared neural mechanisms contributing to psychopathology in both conditions and argue that PIP is a model of more general psychoses.

## Introduction

The overall prevalence of psychosis in people with epilepsy (PWE) is estimated at 5.6%, representing a 7.8-fold increase compared to the general population (Clancy *et al*., 2014). Psychosis in epilepsy is usually classified by its relation to seizure activity: postictal psychosis (PIP) occurs immediately after a seizure or within seven days of return to normal mental function following a seizure, while interictal psychosis is unrelated to seizure activity. PIP occurs in around 2% of PWE (Clancy *et al*., 2014), typically lasts between 24 hours and 3 months (Logsdail and Toone, 1988) and is associated with aggressive behaviours as well as poor epilepsy and psychosocial outcomes (Tarrada *et al*., 2022). Due to its unique presentation (i.e. consequent to a seizure) and modest phenomenological overlay with interictal psychosis (i.e. lack of delusional symptoms or thought disorder), PIP has historically been considered a separate entity to other forms of psychosis with differing underlying mechanisms (Kanemoto *et al*., 1996). However, emerging evidence suggests a shared genetic susceptibility to developing epilepsy and psychosis (Clarke *et al*., 2012, Campbell *et al*., 2020). Braatz *et al*. (2021) showed that PWE who develop PIP have a substantially higher polygenic risk score for schizophrenia than PWE who do not develop PIP. Based on this, they hypothesized that PIP may arise from a genetic susceptibility combined with the additional brain insult of seizures (especially seizure clusters on a background of chronic epilepsy), temporarily exceeding a biological threshold for psychosis. In this study we further investigate whether PIP and schizophrenia share cognitive and electrophysiological features, arguing for PIP to be considered as an organic model for psychoses.

Previous studies have shown that people with chronic schizophrenia, as well as those with early stage psychosis, demonstrate impaired performance at both recognition (Pelletier *et al*., 2005) and associative (or relational) memory tasks (Avery *et al*., 2019), as well as tasks which test the ability to infer associations across memorised pairs that overlap via a shared common item (Armstrong *et al*., 2012; Armstrong, Williams and Heckers, 2012; Armstrong *et al*., 2018; Adams *et al*., 2020). Associative memory function in both healthy cohorts and PWE has been found to depend on theta band activity in medial temporal lobe structures (Klimesch *et al*., 1996; Fell *et al*., 2011; Backus *et al*., 2016; Vivekananda et al., 2020; Joensen et al., 2023; see Herweg *et al*., 2020 for a review). In addition, the inference of memory associations has been shown to depend on theta band functional connectivity between the medial temporal lobe (MTL) and medial prefrontal cortex (mPFC; Backus *et al*., 2016), which is known to be impaired in both rodent models of schizophrenia (Dickerson, Wolff and Bilkey, 2010; Sigurdsson *et al*., 2010) and people with schizophrenia (Adams *et al*., 2020). To our knowledge, however, associative memory function, inference, and associated neural dynamics have not previously been explored in PIP. We aimed to address that shortfall by recording high-density EEG from a group of patients with PIP and matched controls while they performed an associative memory task that probes recognition, associative retrieval, and inference. We hypothesised that people with PIP would show similar behavioural impairments to people with schizophrenia (i.e. poorer recognition, poorer associative memory accuracy, and reduced memory inference); and that this would be associated with deficits in task-related theta power.

## Methods

### Participants

This study was approved by the local NHS ethics committee (REF: 15/LO/1642). We recruited 9 participants with epilepsy and PIP and 9 age matched controls with epilepsy without PIP (see Table 1 for further details). These were drawn from the patient cohort described in Braatz *et al*. (2021) and from clinic lists of patients attending the Chalfont Centre for Epilepsy, Chalfont St Peter, UK. All participants gave written informed consent prior to enrolment in the study and were deemed to have the mental capacity to do so. Our inclusion criteria were: diagnosis of epilepsy, diagnosis of PIP made by a neuropsychiatrist or clearly fulfilling the criteria set by Logsdail and Toone (1988), and the ability to tolerate a 4-5 hour experimental session. Our exclusion criteria were lack of a clear diagnosis of PIP and inability to tolerate the full experimental session. Following Braatz *et al*. (2021) we did not exclude individuals with interictal psychosis, provided that they still experienced clear-cut PIP (although no participants had active psychosis during testing). We recorded handedness, age, gender, phenotypic data regarding epilepsy, psychiatric medical history and current medication of each participant using a standardised proforma. This information was obtained by review of clinical notes and interview of participants at the start of the experimental session. Importantly, no participants showed any evidence of hippocampal sclerosis in MRI, where available.

**Table 1:** Demographics and Clinical Phenotype.

|  | PIP | Control | p-value |
| --- | --- | --- | --- |
| *N* | 9 | 9 |  |
| Left-Handed, *n* | 2 | 0 | 0.4706<sup>F</sup> |
| Age, years; median (IQR) | 40 (24) | 40 (18.75) | 0.8137<sup>MW</sup> |
| Male, *n* | 6 | 2 | 0.1534<sup>F</sup> |
| Focal Epilepsy, *n* | 8 | 6 | 0.5765<sup>F</sup> |
| Known lesion, *n* | 3 | 0 | 0.2088<sup>F</sup> |
| Seizures in last month, *n*; median (IQR) | 3 (7.5) | 3 (9.75) | 0.9999<sup>MW</sup> |
| Concurrent mental health diagnosis (other than PIP), *n* | 4 | 5 | 0.9999<sup>F</sup> |
| Number of different ASMs taken daily, *n* | 3 (1.5) | 2 (1) | 0.5483<sup>MW</sup> |
| Regular use of sodium channel blocking* ASMs, *n* | 5 | 7 | 0.6199<sup>F</sup> |
| Regular or recent (within the last 3 days) use of benzodiazepines, *n* | 3 | 1 | 0.5765<sup>F</sup> |
| Current cannabis usage, *n* | 1 | 1 | 0.9999<sup>F</sup> |
| Current anti-psychotic usage, *n* | 2 | 1 | 0.9999<sup>F</sup> |
| IQR = Interquartile range, ASMs = Anti-seizure medications <sup>F</sup> = Fisher's Exact test, <sup>MW</sup> = Mann-Whitney *U*<br>*Such as Carbamazepine, Lamotrigine, Zonisamide and Phenytoin |  |  |  |

### Memory inference task

Participants undertook a memory inference task similar to that employed by Adams *et al*. (2020), implemented in Psychtoolbox (Brainard, 1997). This task consisted of 3 blocks of ‘study’ and ‘test’ phases during which participants passively viewed pairs of images displayed on screen and then had their memory for those images tested, respectively. Each study trial consisted of a 1 s fixation cross, followed by a pair of images presented for 8 s, followed by a 1 s blank screen (see Figure 1 for further details). Participants were instructed to imagine each pair of images interacting as vividly as possible. Each image was either an object, place or famous person, and each pair of images was drawn from five object-place-person ‘events’ in each block. Crucially, however, only two pairs of images from each event were shown during the study phase, to give a total of 10 image pairs / study trials in each block (and 30 image pairs / study trials overall). The order and content of image pairs was pseudorandomised such that one pair from each event was shown in a random order before the second pair from any event was shown (also in a random order), and the image categories used in the two pairs shown from each event was counterbalanced across events in all blocks (such that participants viewed an equal number of person–place, place–object, and object–person pairs overall).

**Figure 1:**
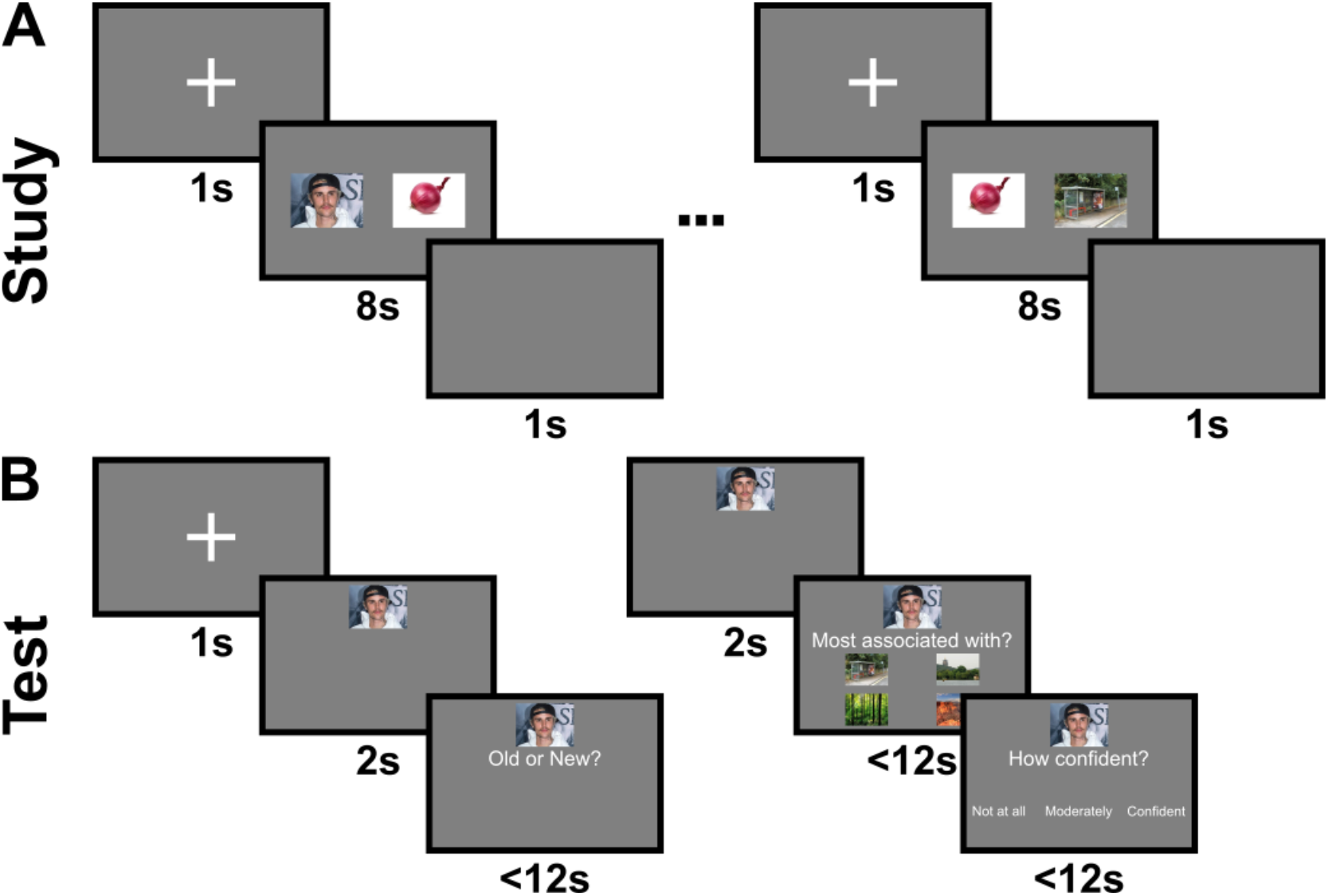
Memory Inference Task. **(A)** During the study phase, participants were shown a series of image pairs for 8 s, preceded by a 1 s fixation cross and followed by a 1 s blank screen. Each image depicted either a person, place, or object. In total, two overlapping image pairs (e.g. Justin Bieber – Onion, Onion - Bus Stop) from five person-place-object events (e.g. Justin Bieber - Onion - Bus Stop) were shown, giving a total of ten study trials in each of three blocks. The third possible pair from each event (e.g. Justin Bieber - Bus Stop) was not displayed but could be inferred. **(B)** During the subsequent test phase, participants were first cued with a single image for 2 s, preceded by a 1 s fixation period, and then asked whether the image shown was ‘old’ (i.e. drawn from one of the image pairs viewed during the study phase) or ‘new’ (i.e. an image that had not been seen before during the task). If the image was ‘old’, regardless of their response, they were then asked to indicate which of four alternative images it was “most associated with” (i.e. had been either directly or indirectly paired with during the study phase). In the example shown here, the inferred association between Justin Bieber and the Bus Stop is being tested. Finally, they were asked to indicate their confidence in that response (‘not at all’, ‘moderately’, ‘confident’). Participants had up to 12 s to respond in each case. All 15 images shown during the study phase were used as a cue in one trial, with all three associations from each event being tested once, alongside an equal number of new items, giving a total of 30 test trials in each of the three blocks.

Each test trial started with a 1 s fixation cross, after which a single ‘cue’ image was displayed for 2 s. Participants were then asked to indicate whether the cue image was ‘old’ or ‘new’ – i.e. whether it had appeared in a pair presented during the study phase or was drawn from an equal number of new events that the participant had not previously seen. If the image was old, regardless of whether the participant responded correctly, there was a further 2 s cue period after which four further images from the same category (i.e. person, place, or object) were presented, one drawn from the same event as the cue and three from other events that had been viewed during the study phase. Participants were asked to indicate which of the four items was “most associated” with the cue image. Importantly, the associations tested were either ‘direct’ (i.e. images that had appeared as a pair during the study phase; two per event, thirty across all blocks) or ‘inferred’ (i.e. images that had appeared in different pairs from the same event, linked via a third image that was common to both pairs; one per event, fifteen across all blocks). Finally, participants were asked to give a confidence rating (‘not at all’, ‘moderately’, or ‘confident’) for their response to the associative memory question. Participants had up to 12 s to answer each memory question and to indicate confidence. All images from all events were used as a cue once during the test phase in each block, and all three associative pairs (two direct and one inferred) from each event were tested once, giving a total of thirty test trials (fifteen for ‘new’ and fifteen for ‘old’ images, of which ten probed direct and five probed inferred associations) in each block, and ninety test trials overall (forty five for ‘new’ and forty five for ‘old’ images, of which thirty probed direct and fifteen probed inferred associations).

Before commencing the main task, participants were shown a set of standard instructions and completed a practice block with the same structure but only two events (i.e. four study trials and twelve test trials, six for old images). Participants had to repeat the practice block until they reached a criterion of >80% performance on recognition memory trials and >50% performance on associative memory trials. All task code, images, and instructions can be found at https://github.com/bushlab-ucl/memoryInferenceTask.

### EEG recording and pre-processing

High-density electroencephalography (EEG) was recorded throughout the task at a sample rate of 5000Hz using 63 actiCAP active electrodes arranged according to the international 10-20 montage in conjunction with an ActiChamp 63-channel amplifier and BrainVision software (Brain Products GmbH, Germany). Electrode impedances were maintained at 5 kΩ or below. EEG recordings were subsequently pre-processed in MATLAB 2019b using the EEGLAB 2021.1 toolbox (Delorme and Makeig, 2004). First, the signal was down-sampled to 200Hz, high-pass filtered at 1Hz to eliminate slow drift and notch filtered in the 49-51Hz band to exclude mains noise. Noisy channels were identified by visual inspection and excluded from all subsequent analyses. The signal on all channels was then re-referenced to a common average reference, and artefacts (i.e. eye blinks, saccades, muscle and channel noise) were visually identified and removed using independent components analysis. Finally, the EEG data were epoched in a [−4 10 s] window around the onset of image pair presentation during each study trial (i.e. from 3s before the onset of the fixation cross to 2s after the end of image pair presentation); and [−4 5 s] window around the onset of each cue presentation during test trials (i.e. from 3s before the onset of the fixation cross to 3s after the end of cue image presentation immediately before the ‘Old or New?’ question was displayed; or 4s before the onset of cue image presentation to 3s after the end of cue image presentation immediately before the ‘Most Associated With?’ question and four alternative forced choice options were displayed; see Figure 1). Trials with excessive noise were identified and excluded from subsequent analyses statistically using the Generalised Extreme Studentised Deviate (GESD) test and by visual inspection. Each recording was inspected for interictal epileptiform discharges but none were found. For our main analyses, this left 25.7 ± 3.83 (mean ± SD, range of 19-30) study trials; 40.3 ± 3.44 (34 – 45) recognition cue trials for ‘old’ items; 39.9 ± 4.89 (30 – 45) recognition cue trials for ‘new’ items; and 37.4 ± 5.29 (29-45) associative cue trials (24.9 ± 3.1 for direct and 12.5 ± 2.53 for inferred pairs) per participant. Importantly, there was no significant difference in the number of study, recognition, or associative cue trials between groups all p>0.22).

### Spectral analysis

Spectral analyses of the pre-processed EEG data were subsequently conducted using the Fieldtrip toolbox (Oostenveld *et al*., 2011). First, time-varying estimates of oscillatory power were obtained using a five cycle Morlet wavelet transform. Next, power estimates in each trial were log transformed and baseline corrected by subtracting the mean power in each frequency band during a [−1 −0.5 s] window before image pair presentation (during study trials) or cue presentation (during test trials). Finally, baseline corrected time-frequency representations were averaged across all trials in each condition of interest. Averaged power values in each time and frequency window of interest were then entered into second level statistical tests to look for differences between groups and / or conditions.

### Source localisation

For source localisation, head models were created based on individual structural MRI scans, where those were available (for five patients in the PIP group and two in the control group). The MRI scans were segmented into brain, CSF, skull, and scalp using SPM12 and the Huang toolbox (Huang *et al*., 2013). A volume conduction model was constructed using the mesh created from the segmented MRI in Fieldtrip, and template electrode positions were aligned to individual head models. The MNI template MRI and a template volume conduction model were used for the remaining individuals, together with re-aligned template electrode positions to the template head model. For group analyses, each individual MRI was warped to a template MRI in Fieldtrip. The inverse of the warp was applied to a template dipole grid so that each grid position was consistent across participants in MNI space. Finally, the raw data were low pass filtered at 30 Hz and baseline corrected using the mean amplitude in a [−1 −0.05 s] window prior to stimulus onset in each trial. A covariance matrix was computed on the data from each epoch, and spatial filters were computed using a linear constrained minimum variance (LCMV) beamformer with 7% regularisation.

### Statistical analysis

For analyses of behaviour, where data was missing for one or more participants in any one condition (because no errors were made, for example), linear mixed-effects models were constructed using RStudio and lme4 (*Bates et al*., 2015). For analyses of the EEG data, cluster-based permutation tests were implemented in Fieldtrip to identify differences between conditions or groups while accounting for the multiple comparisons problem (Maris & Oostenveld, 2007). Our analyses focussed on the 3-7 Hz theta band identified by previous studies of memory inference in healthy participants (Backus *et al*., 2016). Power values in this frequency band were averaged across the entire 8 s image pair presentation time window (for the study phase) or entire 2 s cue image presentation time windows (for the test phase) to provide a single measure of average theta power for each EEG channel or source space voxel. These power values were then entered into cluster-based permutation tests using the Monte Carlo method with either a two-sided dependent-samples t-test (for within group analyses) or independent-samples t-test (for between group analyses). The minimum number of neighbouring channels / voxels within a spatial cluster was three / six (at the sensor and source level, respectively), with neighbouring channels / voxels defined based on the triangulation method / connectivity in a 3d volume. In each case, 5000 random permutations of condition labels (for within group analyses) or group labels (for between group analyses) were performed, t-statistic values were summed across groups of neighbouring channels / voxels that showed significant effects at an uncorrected threshold of p<0.05 to form cluster-level statistics, the maxima of which were recorded for each randomisation. Cluster-corrected p-values were then calculated as the proportion of cluster-level statistics across all randomisations that were larger than the true cluster-level statistics in the data.

## Results

We recorded high density scalp EEG from nine participants with both epilepsy and post-ictal psychosis (PIP) and nine matched control participants with epilepsy (PWE) but without PIP while they completed a memory inference task that probed recognition memory, associative memory, and the inference of unseen associations – each of which is known to be impaired in schizophrenia (Scz; see Methods for further details). Previous studies have demonstrated that both associative memory accuracy and inference can be predicted by changes in frontotemporal theta power and functional connectivity (Klimesch *et al*., 1996; Fell *et al*., 2011; Backus *et al*., 2016; Herweg, Solomon and Kahana, 2020; Joensen *et al*., 2023). Our aim was to establish whether the same pattern of behavioural deficits was present in PIP, and whether this was associated with disrupted theta dynamics.

### Associative Memory Accuracy and Memory Inference are Impaired in PIP

First, to quantify recognition memory performance, we computed the sensitivity index (d’) and response bias for each participant. Comparing across groups, we found no significant differences in d’, which was high in both PIP and control groups (median values of 3.71 and 3.99, respectively; *p* = 0.229, Mann-Whitney U test). There was also no significant difference in response bias across groups (*p* = 0.881, independent samples t-test). Consistent with this we found no significant difference in the rate of hits (median value of 97.8% in both groups; *p* = 0.613, Mann-Whitney U test) or correct rejections (median value of 97.8% in PIP and 100% in controls; *p* = 0.367, Mann-Whitney U test) between groups (Figure 2A). This is in contrast to people with Scz who tend to show impaired recognition memory compared to healthy controls (Pelletier *et al*., 2005) as well an increased tendency towards false alarms (Adams *et al*., 2020), although subtle differences may have been missed as performance was near ceiling in both groups.

**Figure 2:**
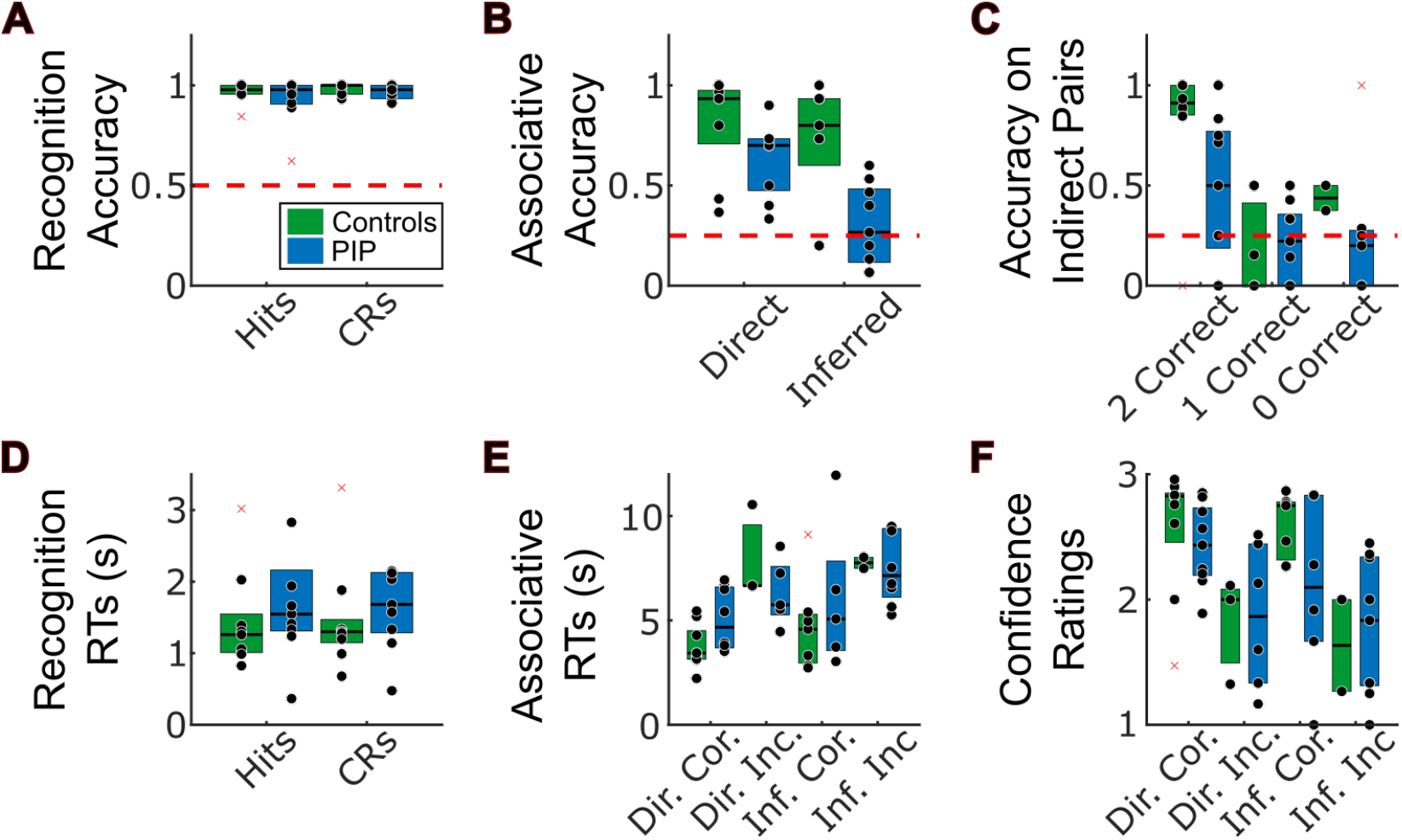
Behavioural Performance. **(A)** Recognition memory accuracy by group for old (hits) and new items (correct rejections, CRs). We found no significant differences between groups in recognition memory performance, **(B)** Associative memory accuracy by group for direct (observed) and inferred (unobserved) pairs. Across both groups, performance on direct associations is significantly better than on inferred associations. In addition, the control group performed significantly better than the PIP group overall. Finally, the difference in memory accuracy between direct and inferred pairs was greater in the PIP group. Notably, the PIP group also performed no better than chance (25%) on inferred pairs. **(C)** Memory inference accuracy split by the number of correctly retrieved direct associations from the same event. Accuracy for inferred pairs is significantly greater than chance in the control group, but not the PIP group, when both direct pairs from the same event were correctly retrieved (2 Correct), but not in either group when either one (1 Correct) or zero (0 Correct) direct pairs from the same event were correctly retrieved. **(D)** Recognition memory reaction times (RTs) by group for old (hits) and new items (correct rejections, CRs). We found no significant differences between groups. **(E)** Associative memory reaction times (RTs) by group for direct (observed) and inferred (unobserved) pairs where responses were correct or incorrect. RTs were faster across all participants for direct pairs and correct responses. **(F)** Confidence ratings for associative memory judgements by group for direct (observed) and inferred (unobserved) pairs where responses were correct or incorrect. Confidence ratings were higher across all participants for direct pairs and correct responses.

Next, we compared associative memory accuracy between groups using a repeated measures ANOVA with a between subject factor of group and within subject factor of direct versus inferred association (Figure 2B). This revealed a main effect of pair type (*F*(1,16) = 22.0, *p* < 0.001), a main effect of group (*F*(1,16) = 8.52, *p* = 0.01), and a pair type x group interaction (*F*(1,16) = 5.97, *p* = 0.026). Post-hoc testing indicated that these results were driven by better memory for direct vs inferred pairs across groups (*t*(17) = 4.12, *p* < 0.001), better associative memory in the control group vs PIP (*t*(16) = 2.92, *p* = 0.01), and the difference between performance on direct and inferred associations being smaller in the control group vs PIP (*t*(16) = −2.44, *p* = 0.027). In addition, memory for both direct and inferred pairs was better than chance in the control group (*t*(8) = 6.94, *p* < 0.001 and *t*(8) = 4.55, *p* = 0.0019, respectively); while for the PIP group only memory for direct pairs was better than chance (*t*(8) = 6.30, *p* < 0.001 and *t*(8) = 0.79, *p* = 0.45, respectively). This reduction in overall associative memory accuracy, as well as the pronounced impairment of inference memory specifically is consistent with the Scz phenotype described by previous studies (Armstrong *et al*., 2012, 2018; Armstrong, Williams and Heckers, 2012; Avery *et al*., 2019; Adams *et al*., 2020).

Next, we aimed to establish whether the marked impairment of inference within the PIP group arose from their reduced memory for direct associations, as participants could only be expected to accurately infer associations from events in which they accurately recalled both linking pairs. To do so, we isolated the inferred associations from events where both linking pairs (direct associations) had been correctly recalled (Figure 2C). In these pairs, the likelihood of a correct inference was also greater than chance for the control group but not the PIP group (*t*(8) = 3.43, *p* < 0.01 and *t*(8) = 1.85, *p* = 0.10, respectively; Figure 2B). In contrast, for inferred associations where one or no direct pairs from the corresponding event were correctly retrieved, neither group performed above chance (all *p* > 0.204).

To further explore this effect, while accounting for the fact that some participants had no instances where zero, one or two direct associations from a given event were accurately retrieved, we constructed a linear mixed-effects model of accuracy on inferred associations with a random factor of participant ID and fixed factors including the number of correct direct associations from the same event and participant group. A likelihood-ratio test suggested a significant effect of the number of correct direct associations (*χ*^2^(1) = 21.93, *p* < 0.001), and examination of the full model indicated that if two direct associations from an event were correctly retrieved, then 38% more inferred associations were correctly retrieved, on average, than when only one or no direct associations were correctly retrieved. In addition, we found a trend towards a significant interaction between group and the number of correct direct associations (*χ*^2^(1) = 3.93, *p* = 0.056), suggesting that this effect was larger in the control group than PIP. In sum, these results suggest that the strongest determinant of inference performance in both groups is associative memory accuracy for the direct pairs in the same event, although there also appears to be a meaningful reduction in inference performance associated with having PIP compared to having epilepsy alone.

Next, we examined reaction times during recognition and associative memory judgements. First, a repeated measures ANOVA with a between subject factor of group and within subject factor of hits vs correct rejections revealed no main effects or interactions (all *p* > 0.315; Figure 2D). Second, a linear mixed-effects model of reaction times during associative memory judgements with a random factor of participant ID and fixed factors including accuracy, pair type (direct vs inferred), and group revealed significant main effects of accuracy (*χ*^2^(1) = 6.26, *p* = 0.017) and pair type (*χ*^2^(1) = 10.66, *p* = 0.002). An examination of the full model indicated that responses were 1.75 s faster for correct pairs and 2.80 s faster for direct associations, on average, across all participants (Figure 2E). No other main effects or interactions were found. Finally, we examined the confidence ratings given by participants following associative memory judgements using a similar linear mixed-effects model with a random factor of participant ID and fixed factors including accuracy, pair type (direct vs inferred), and group. Again, this revealed significant main effects of accuracy (*χ*^2^(1) = 5.85, *p* = 0.021) and pair type (*χ*^2^(1) = 16.57, *p* < 0.001), with confidence ratings being 0.19 higher for correct pairs and 0.40 higher for direct associations, on average (Figure 2F). No other main effects or interactions were found.

### Increased Frontotemporal Theta Power during Successful Memory Encoding in Controls but not PIP

Successful associative memory formation can typically be predicted by increased theta band oscillatory power in medial temporal lobe regions including the hippocampus, both in healthy cohorts and people with epilepsy (Klimesch *et al*., 1996; Fell *et al*., 2011; Joensen *et al*., 2023; see Herweg *et al*., 2020 for a review). Hence, we next examined changes in oscillatory power during the 8 s study period, averaged across all channels and participants (Figure 3A). This revealed an increase in low frequency power that peaked shortly after the onset of image pair presentation and persisted throughout the 8 s study period. We then compared oscillatory power between the presentation of image pairs that were and were not subsequently remembered. This identified a more pronounced increase in low frequency power for subsequently remembered image pairs in the control group than in the PIP group (Figure 3B).

**Figure 3:**
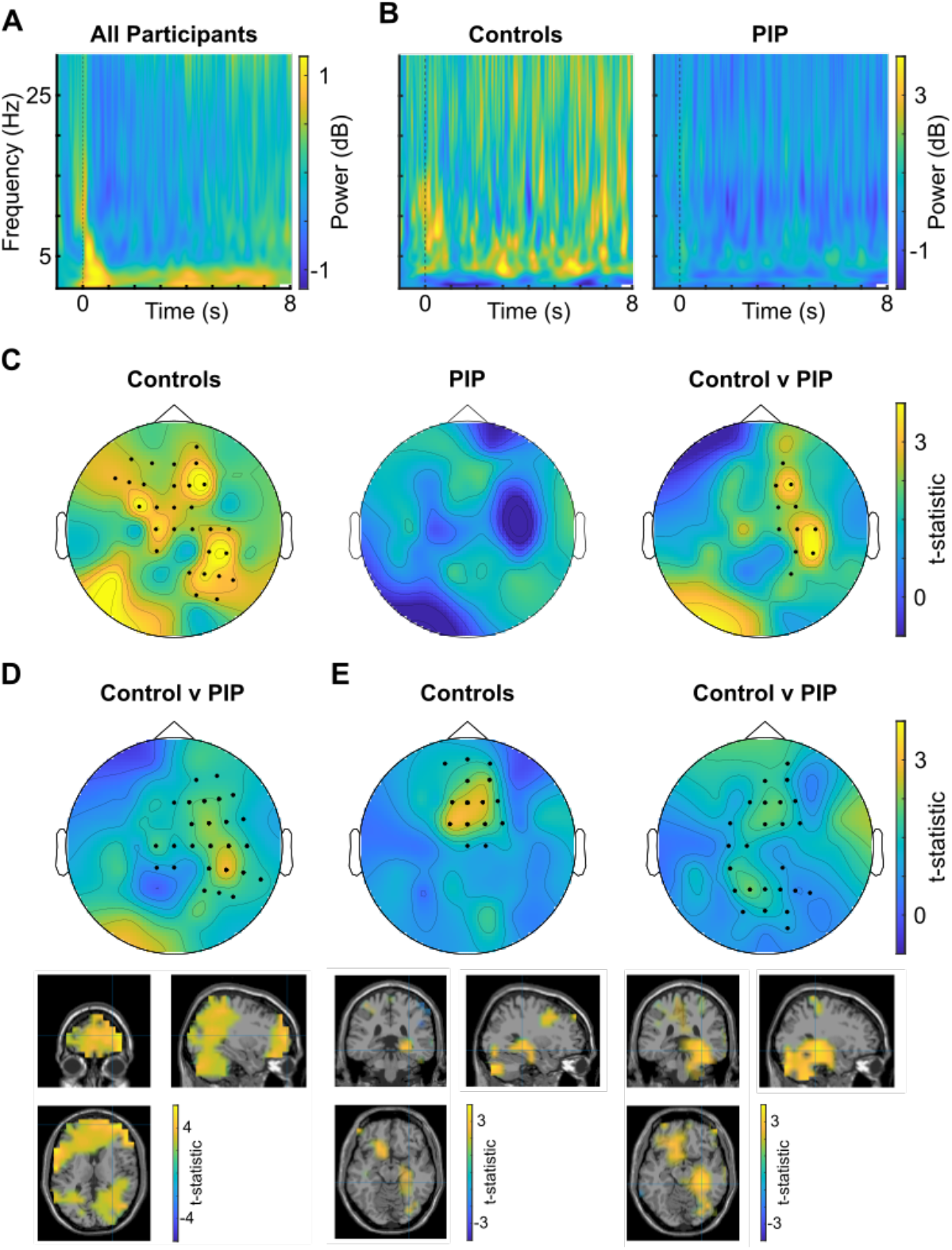
Greater 3-7 Hz theta power during successful memory formation in controls, but not PIP. **(A)** Baseline corrected time-frequency plot of the 8 s encoding period, averaged across all sensors and both groups. **(B)** Baseline corrected time-frequency plots of the subsequent memory effect (i.e. correct – incorrectly remembered pairs) over the 8 s encoding period, averaged across all sensors and shown for the control group (left) and PIP group (right) separately. **(C)** Scalp map of 3-7 Hz theta power subsequent memory effect across the 8 s encoding period for the control group (left), PIP group (middle), and for the difference between groups (right). This shows that the control group have significantly greater right frontotemporal theta power for subsequently remembered vs forgotten pairs during encoding, compared to the PIP group. **(D)** Increased 3-7 Hz theta power for subsequently remembered vs forgotten pairs during the 2.5-3.65 s time window identified by cluster analysis is significantly stronger in controls vs the PIP group. Importantly, this effect passes our statistical threshold for cluster level correction in *a priori* bilateral hippocampus and mPFC ROIs. **(E)** Cluster analysis indicates that the control group also show significantly stronger 3-7 Hz theta power over frontal midline regions during the encoding of pairs in events where the inference judgement is subsequently made correctly (top left), and that this effect is significantly stronger in the control vs PIP group (top right). This subsequent inference effect can be source localised to the right medial temporal lobe (MTL) in the control group (bottom left) and in the contrast between groups (bottom right). In each case, these effects pass out statistical threshold for cluster level correction in an *a priori* bilateral hippocampus ROI. In panels C-E, highlighted sensors pass our cluster level significance threshold of *p* < 0.05.

To further probe this effect, we focussed our analyses on a 3-7 Hz theta frequency band that has been previously implicated in associative memory and memory inference function during encoding (Backus *et al*., 2016). First, we averaged 3-7 Hz theta power across the whole 8 s window and across all EEG channels, then used a 2 x 2 repeated measures ANOVA to look for differences between subsequently remembered vs subsequently forgotten pairs in participants from the control vs PIP group. This revealed a main effect of subsequent memory performance (*F*(1,14) = 18.5, *p* < 0.001, 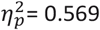) on theta power, as well as a significant interaction between subsequent memory and group (*F*(1,14) = 8.05, *p* = 0.013 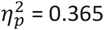). Post-hoc tests indicated that this interaction was driven by a subsequent memory effect in the control group (*t* = 4.16, *p* = 0.006, Cohen’s *d* = 1.57) but not in the PIP group (*t* = 1.26, *p* = 0.244, Cohen’s *d* = 0.419).

Next, to identify memory related theta oscillations at the sensor-level, we averaged baseline corrected 3-7 Hz power across the whole 8 s study period for each EEG channel and compared subsequently remembered and forgotten pairs (Figure 3C). Consistent with the results above, this revealed a large, right lateralised frontotemporal cluster in the control group (cluster *t* = 89.6, *p* < 0.001) but no significant cluster in the PIP group, with a significant difference between groups in the same region (cluster *t* = 31.3, *p* = 0.016). These results suggest that successful associative memory formation is associated with increased theta power over frontotemporal regions in control participants, consistent with previous studies, but not in those with PIP.

Next, we used permutation statistics to identify specific time windows within the 8 s study period during which average 3-7 Hz theta power across all sensors was significantly greater during the presentation of image pairs that were subsequently remembered vs forgotten in controls vs PIP. This revealed two periods, from 2.5-3.65 s and 5.3-5.8 s after the onset of image pair presentation. We focussed on the first (longer) time window and attempted to localise the source of associative memory related theta power during this period. This revealed a significant cluster in the right frontal cortex in which 3-7 Hz theta power was greater for subsequently remembered vs forgotten image pairs in the control group than the PIP group (cluster *t* = 2720.5, *p* < 0.001; Figure 3D). Importantly, this theta power subsequent memory effect was also significant for the control group in *a priori* defined bilateral hippocampal (cluster *t* = 3.042, *p* = 0.016) and medial prefrontal (cluster *t* = 353.6, *p* < 0.001) ROIs during the same time window.

Finally, we focussed on the encoding of image pairs (A-B, B-C) from events where the unobserved pair (A-C) was correctly inferred during retrieval, compared to those where this inference was not correctly made. Spatiotemporal permutation analysis identified that 3-7 Hz theta power in the control group was significantly greater over frontal midline sensors from 1.95-3s after the presentation of image pairs from events where accurate inference was subsequently made (cluster *t* = 365.9, *p* = 0.034; Figure 3E). Moreover, the difference in 3-7 Hz theta power between pairs from events where the inferred pair was accurately vs not accurately identified during this time window was also significantly stronger in the control group vs PIP (cluster *t* = 22.8, *p* = 0.018; Figure 3E). Source localisation indicated that this inference related theta power in the control group originated from right medial temporal lobe (MTL) regions during the same time window, as did the difference between groups. In both cases, this theta power subsequent inference effect was significant in an *a priori* defined bilateral hippocampal ROI during this time window (control group: cluster *t* = 5.97, *p* = 0.024; group difference: cluster *t* = 12.9, *p* < 0.001). Overall, these findings indicate that MTL theta power is greater during the encoding of image pairs from events that support subsequent inference in controls, consistent with previous studies (Backus *et al*., 2016), but not in participants with postictal psychosis.

### Greater Frontotemporal Theta Phase Coupling in the Control Group

In addition to increased hippocampal theta power, previous studies have reported that increased theta band functional connectivity between frontal and medial temporal lobe regions during memory encoding supports later inference (Backus *et al*., 2016). Hence, we next examined changes in 3-7 Hz theta phase coupling during encoding, between the mPFC region that showed the greatest theta subsequent memory effect across all participants (Figure 4A) and an *a priori* defined right hippocampal ROI, which showed increased theta power during the encoding of image pairs from events where the unobserved pair was later correctly inferred (Figure 3E). First, we averaged 3-7 Hz theta phase coupling between these regions across the whole 8 s window, then used a 2 x 2 repeated measures ANOVA to look for differences between image pairs from subsequently inferred vs not inferred events in participants from the control vs PIP group (Figure 4B). This revealed a significant interaction between group and subsequent inference (*F*(1,13) = 7.13, *p* = 0.019), but no main effects (both *p* > 0.21). Post-hoc testing indicated that this was driven by and a greater difference between theta phase coupling during the encoding of pairs that later supported accurate inference in the control group vs PIP (*t*(13) = 2.67, *p* = 0.0193). Importantly, this arose from a trend towards greater frontotemporal theta phase coupling for subsequently inferred events in the control group (*t*(6) = 2.29, *p* = 0.0622), with no clear difference in the PIP group (*t*(7) = −1.25, *p* = 0.252).

**Figure 4:**
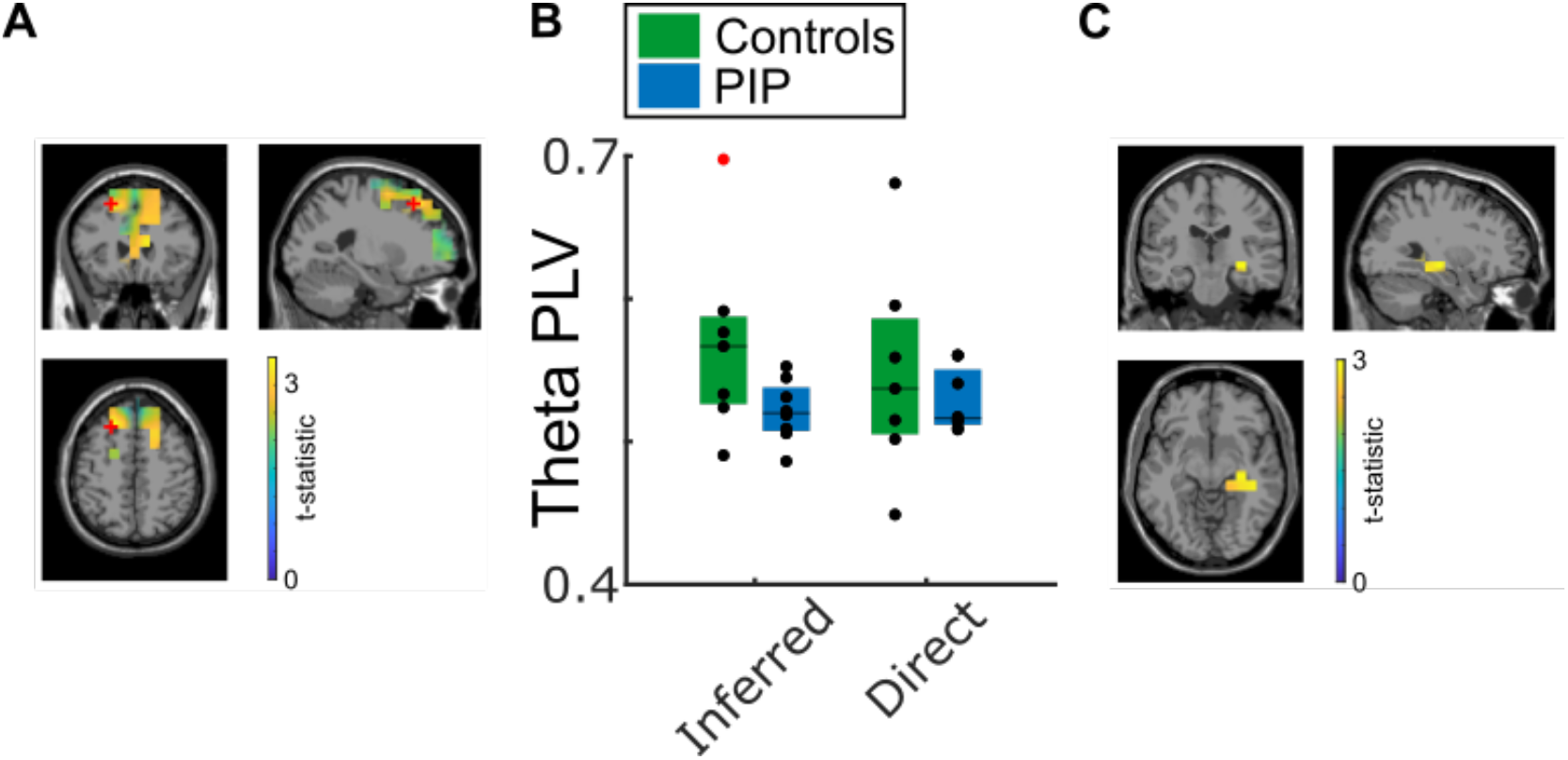
Subsequent inference is supported by 3-7 Hz frontotemporal theta phase coupling during memory encoding. **(A)** Source localisation of 3-7 Hz theta power for subsequently remembered vs forgotten image pairs during the 8 s encoding period, averaged across all sensors and both groups. The peak mPFC voxel from this contrast (marked with a red cross) is used as a seed to examine theta phase coupling with an *a priori* defined right hippocampal ROI. **(B)** mPFC theta phase locking values averaged across all voxels within an *a priori* defined right hippocampal ROI, during the encoding of image pairs from events where the unobserved pair was later inferred vs not inferred in the control and PIP groups. The difference in theta phase coupling between subsequently inferred and not inferred events is significantly greater in controls vs PIP. **(C)** Source localisation of the 3-7 Hz theta phase coupling subsequent inference effect across the 8 s encoding period for the control group. This shows that the control group have significantly greater theta phase coupling between mPFC and a cluster in the right hippocampus for subsequently inferred vs not inferred events during encoding.

Consistent with this, source localisation revealed two significant clusters in the posterior right hippocampus that showed greater theta phase coupling with the mPFC for subsequently inferred vs not inferred events in the control group vs PIP (cluster *t* = 10.69, *p* = 0.004 and cluster *t* = 3.25, *p* = 0.03, respectively). Moreover, examining theta phase coupling in each group separately revealed a right posterior hippocampal cluster in the control group that showed significantly greater theta phase coupling with our mPFC seed for subsequently inferred events (cluster *t* = 8.97, *p* = 0.03; Figure 4C). In contrast, the PIP group showed a significant cluster in the same region that showed reduced theta phase coupling with the mPFC seed for subsequently inferred events (cluster *t* = −2.68, *p* = 0.04; data not shown). These results suggest that increased theta band functional connectivity between frontal and right medial temporal lobe regions supports subsequent memory inference in our control participants but not those with PIP, consistent with previous studies (Backus *et al*., 2016).

### No Difference Between Groups in Theta Power during Recognition Memory Judgements

Next, we examined the neural correlates of successful recognition memory. First, we computed average oscillatory power across all channels and participants during the cue period immediately preceding the old vs new recognition memory judgement. This revealed an increase in low frequency power from baseline that persisted throughout this 2 s recognition cue period (Figure 5A). Next, we compared low frequency power during ‘hit’ trials (i.e. correctly recognised cue images from ‘old’ events) with that during ‘correct rejection’ trials (i.e. correctly recognised cue images that had not formed part of any observed pair). In this case, however, there was no clear difference in oscillatory dynamics between control participants and those with PIP (Figure 5B).

**Figure 5:**
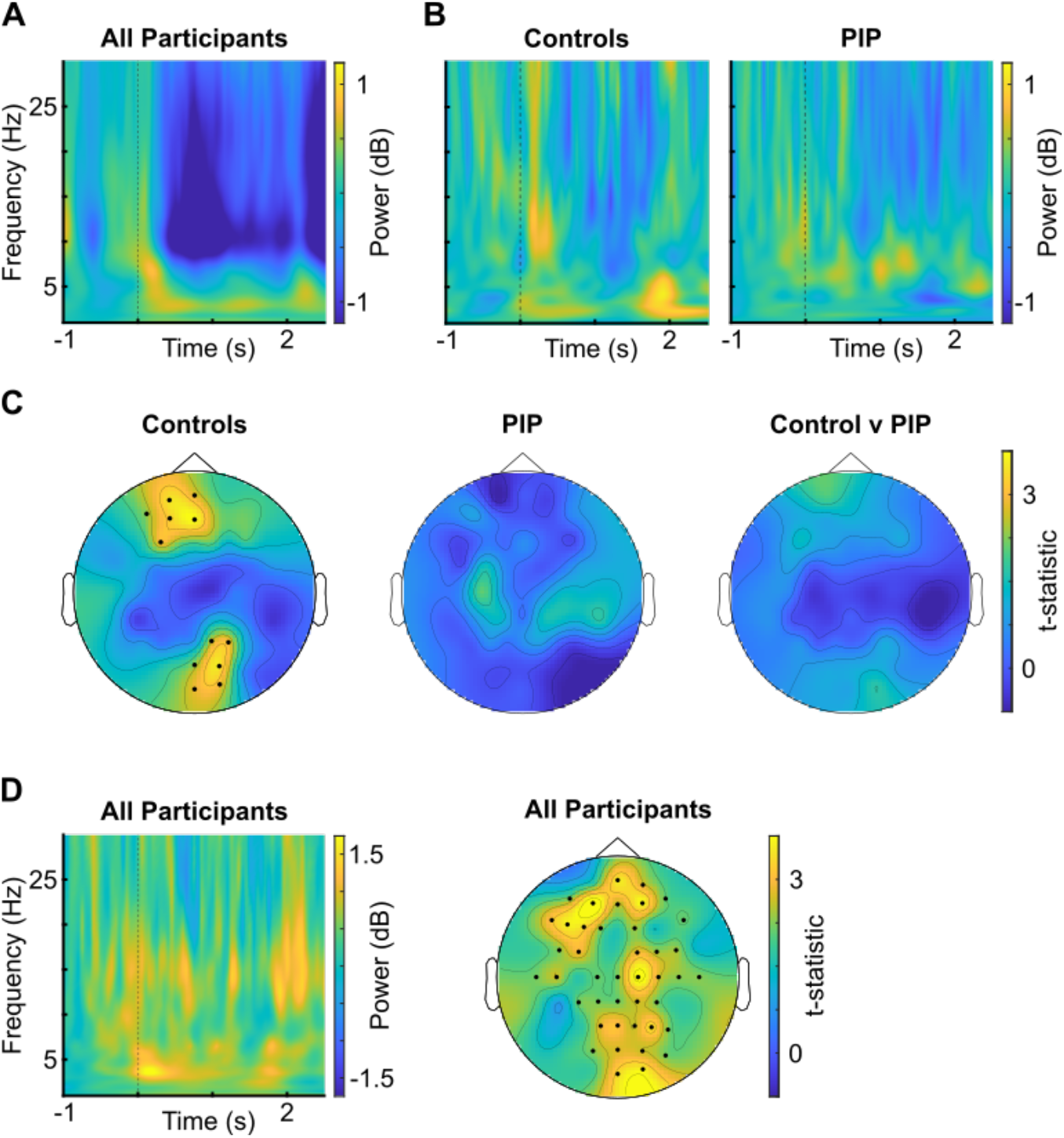
No difference in 3-7 Hz theta power during recognition memory judgements between groups. **(A)** Baseline corrected time-frequency plot of the 2 s recognition cue period, averaged across all sensors and both groups. **(B)** Baseline corrected time-frequency plots of hits vs correct rejections over the 2 s recognition cue period, averaged across all sensors and shown for the control group (left) and PIP group (right) separately. **(C)** Scalp map of 3-7 Hz theta power for hits vs correct rejections across the 2 s recognition cue period for the control group (left), PIP group (middle), and for the difference between groups (right). Although only the control group exhibit significant clusters of increased theta power over frontal midline and occipital regions, there is no significant difference between groups. **(D)** Baseline corrected time-frequency plot of accurate vs inaccurate associative retrieval over the 2 s recognition cue period, averaged across all sensors and both groups (left); and scalp map of 3-7 Hz theta power for accurate vs inaccurate associative retrieval over the 2 s recognition cue period for all participants (right).

To quantify this, we averaged 3-7 Hz theta power across the entire 2 s recognition cue period and all EEG channels and entered those values into a 2 x 2 repeated measures ANOVA with factors of trial type (hits vs correct rejections) and group (control vs PIP). This revealed a trend towards significance for a main effect of trial type (*F*(1,16) = 3.96, *p* = 0.064, 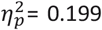), but no other main effects or interactions. This was corroborated by an examination of 3-7 Hz theta power across sensors, which revealed two significant clusters in control participants (anterior cluster *t* = 19.7, *p* = 0.015; posterior cluster *t* = 18.1, *p* = 0.021) where theta power was greater during the correct identification of old vs new items (i.e. for ‘hits’ vs ‘correct rejections’), but none in PWE with PIP and no significant differences between groups (Figure 5C). Overall, these results suggest an increase in theta power during the accurate recognition of old images, compared to new images, which does not reach significance and does not differ between control participants and those with PIP, consistent with previous studies (Osipova *et al*., 2006).

Finally, we asked whether theta power during the recognition cue period might also predict later associative memory retrieval success. To quantify this, we entered average 3-7 Hz theta power values during the recognition cue period of ‘old’ events into a 2 x 2 repeated measures ANOVA with factors of group (control vs PIP) and subsequent associative retrieval accuracy (correct vs incorrect). This revealed a significant main effect of retrieval accuracy (*F*(1, 15) = 23.9, p < 0.001, 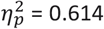) but no other main effects or interactions. This was corroborated by an examination of 3-7 Hz theta power across sensors for all participants, which revealed a large midline cluster where power was significantly greater preceding accurate vs inaccurate associative retrieval (cluster *t* = 119.7, *p* < 0.001; Figure 5D). Hence, when presented with a recognition cue image that was observed in one or more pairs during encoding, theta power is greater when those pairs are subsequently retrieved in both our control group and PWE with PIP.

### Increased Theta Power in the Temporal Lobe during Associative Memory Inference in Controls but not PIP

In addition to theta subsequent memory effects during encoding, previous studies have indicated increases in frontotemporal theta power in healthy cohorts and people with epilepsy during successful associative memory retrieval (Kaplan *et al*., 2014; Vivekananda *et al*., 2021; see Herweg *et al*., 2020 for a review). Hence, we next examined changes in oscillatory power during cued associative memory retrieval and the time window immediately afterwards, where participants were asked to identify the associated image (see Methods and Figure 1 for further details). This revealed increases in low frequency power throughout this time window that peaked shortly after the start of the ‘associative choice’ period, when the four alternative forced choice images were displayed (Figure 6A).

**Figure 6:**
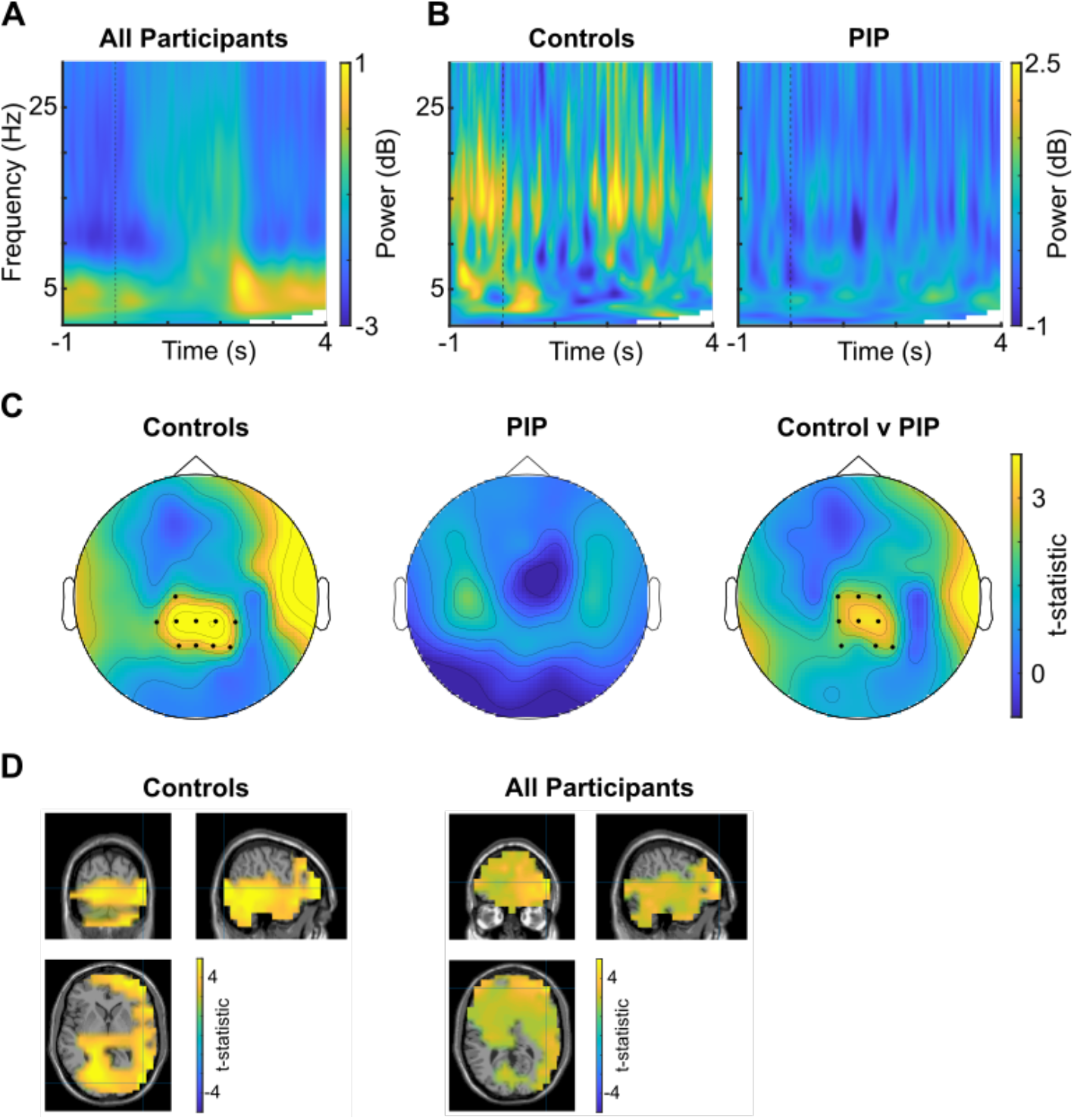
No difference in 3-7 Hz theta power during associative memory judgements between groups. **(A)** Baseline corrected time-frequency plot of the 2 s associative retrieval cue and start of the subsequent associative choice period, averaged across all sensors and participants. **(B)** Time-frequency plot of the 2 s associative retrieval cue and start of the subsequent associative choice period, baseline corrected by data from the fixation period, for accurate vs inaccurate associative memory retrieval, for the control (left) and PIP (right) groups. **(C)** Scalp map of 3-7 Hz theta power during the first 2 s of the associative choice period for accurate vs inaccurate associative memory retrieval in the control (left) and PIP groups (middle), and for the contrast between groups (right). **(D)** Source localisation of 3-7 Hz theta power during the first 2 s of the associative choice period for accurate vs inaccurate associative memory retrieval in the control group (left) and across all participants (right). In each case, this effect passes our statistical threshold for cluster level correction within *a priori* bilateral hippocampus and bilateral mPFC ROIs. In panel C, highlighted sensors pass our cluster level significance threshold of p < 0.05.

Next, we compared low frequency power between cue periods for ‘old’ items preceding accurate versus inaccurate associative retrieval. This revealed increased theta power preceding accurate retrieval that peaked shortly after the onset of the cue image and again after the onset of the associative choice period, which appeared to be more prominent in the control group (Figure 6B). To quantify this, we averaged 3-7 Hz power across all channels and the whole 0-2s associative cue period, baseline corrected by data from the fixation period, and entered those values into a 2×2 ANOVA with factors of correct vs incorrect retrieval and control vs PIP. This revealed a main effect of accuracy, regardless of group: *F*(1, 15) = 4.671, *p* = 0.047, 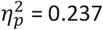; but no other main effects or interactions.

Next, we conducted the same repeated measures ANOVA on average 3-7 Hz theta power during the first 2 s of the associative choice period, which immediately followed the associative retrieval cue, baseline corrected by average power during the immediately preceding associative cue period. This revealed a significant main effect of retrieval accuracy (*F*(1,15) = 5.50, *p* = 0.033, 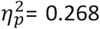) and a trend towards an interaction between retrieval accuracy and group that did not reach significance (*F*(1,15) = 3.27, *p* = 0.091, 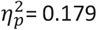). This was corroborated by an examination of 3-7 Hz theta power across sensors, which revealed a significant posterior central cluster in controls (cluster *t* = 35.1, *p* = 0.004), no significant clusters in PIP, and a significant difference between groups in the same region (cluster *t* = 24.2, *p* = 0.018; Figure 6C).

Source localisation suggested that increases in theta power during the first 2 s of the associative choice period preceding accurate memory retrieval in both the control group and across all participants originated from a widespread, right lateralised network of regions with peaks in frontal and occipital cortex (Figure 6D), while there were no significant clusters in the PIP group (data not shown). Importantly, both for the control group and across all participants, theta power during this period was significantly greater for accurate associative memory retrieval in both an *a priori* defined bilateral hippocampal (controls: cluster *t* = 13.0, *p* = 0.004; all participants: cluster *t* = 11.8, *p* = 0.004) and medial prefrontal (controls: cluster *t* = 222.7, *p* = 0.004; all participants: cluster *t* = 344.1, *p* < 0.001) ROIs during the same time window.

Finally, we asked whether there were any differences in theta power between the retrieval of inferred vs direct associations during the same time window. To do so, we first contrasted low frequency power between the successful retrieval of inferred vs direct pairs in each group, averaged across all sensors. This showed an increase in theta power during successful inference that peaked shortly after the onset of the associative cue period, and again after the onset of the associative choice period, in the control group but not in PIP (Figure 7A). To quantify this, we averaged 3-7 Hz power across all channels and the whole 0-2 s associative cue period, baseline corrected by data from the fixation period, and entered those values into a 2×2 ANOVA with factors of accurate inferred vs direct retrieval and control vs PIP. However, this revealed no main effects or interactions (all *p* > 0.47). Next, we carried out the same analysis on average 3-7 Hz theta power from the 2-4 s associative choice period. This revealed a main effect of group (*F*(1,16) = 9.94, *p* = 0.006, 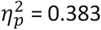) and a significant interaction (*F*(1,16) = 6.23, *p* = 0.024, 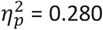). Post-hoc tests revealed that this interaction was driven by theta power being greater during the accurate retrieval of inferred vs direct pairs in controls, but not in PIP (*t*(16) = 2.50, *p* = 0.024, Cohen’s d = 1.18). This was corroborated by an inspection of time-frequency plots, averaged across all sensors, and contrasted between groups (Figure 7B).

**Figure 7:**
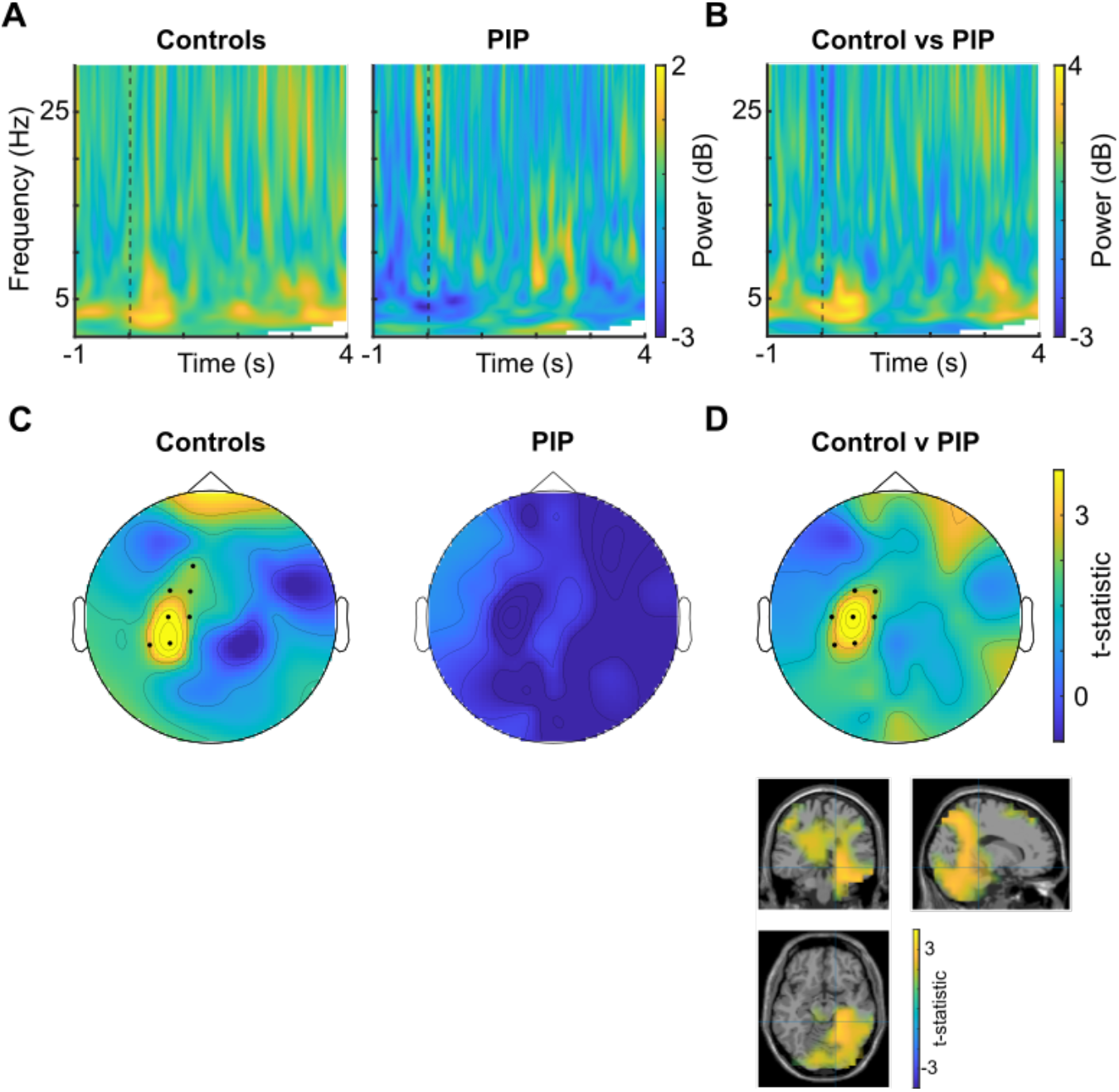
Greater 3-7 Hz theta power during successful memory inference in controls, but not PIP. **(A)** Baseline corrected time-frequency plot of the 2 s associative retrieval cue and start of the subsequent associative choice period, for accurate inferred vs direct associative memory retrieval, for the control (left) and PIP (right) groups. **(B)** Baseline corrected time-frequency plot of the 2 s associative retrieval cue and start of the subsequent associative choice period, for accurate inferred vs direct associative memory retrieval, compared between groups. **(C)** Scalp map of 3-7 Hz theta power during the first 2 s of the associative choice period for accurate inferred vs direct associative memory retrieval in the control (left) and PIP (right) groups. **(D)** Scalp map (top) and source localisation (bottom) of 3-7 Hz theta power during the first 2 s of the associative choice period for accurate inferred vs direct associative memory retrieval, compared between groups.

Next, we examined the sensor-level origin of inference related theta power. In the control group, we found a significant cluster over left temporal regions that showed greater theta power during the retrieval of inferred vs direct associations (cluster *t* = 23.2, *p* = 0.006); while no such clusters were identified in the PIP group (Figure 7C). Crucially, this increase in theta power over left temporal sensors during the retrieval of inferred associations was also significantly greater in the control group vs PIP (cluster *t* = 21.6, *p* = 0.017; Figure 7D), and source localisation indicated that this effect originated from similar right posterior temporal lobe regions as the associative memory effect described above. Again, this effect was significantly greater for the control group vs PIP in the *a priori* defined right hippocampal ROI during this time window (cluster *t* = 8.42, *p* = 0.014).

## Discussion

Our behavioural data show impaired associative memory and memory inference in participants with PIP compared to participants with epilepsy but no history of PIP. This pattern of impairment is identical to that previously described in schizophrenia and early phase psychosis (Armstrong *et al*., 2012, 2018; Armstrong, Williams and Heckers, 2012; Avery *et al*., 2019; Adams *et al*., 2020), suggesting a mechanistic link between these conditions. In particular, our PIP cohort exhibited greater impairment of memory inference than the retrieval of directly observed associations, to the extent that memory inference was no better than chance even when both direct associations were correctly remembered. In contrast, we found no differences in recognition memory performance between groups, in contrast to previous studies which have demonstrated that schizophrenia is associated with impaired recognition memory (Pelletier *et al*., 2005) and an increased tendency towards false alarms (Adams *et al*., 2020). However, we note that recognition memory performance was close to ceiling in our study, as the task was simplified for clinical participants, which may have limited our ability to detect such differences. In addition, the sample sizes for both groups were small (n=9). However, the reliability of our findings is bolstered by a well-matched control group of PWE but without psychosis, a strongly hypothesis-led approach, and group differences of large effect size.

Our neuroimaging data demonstrate that the theta power increases typically associated with successful memory encoding in healthy controls and people with epilepsy were also absent in the PIP group. Specifically, our control participants – but not those with PIP – showed greater theta power during the encoding of image pairs that were later retrieved. This effect appeared to originate from frontal, temporal, and occipital regions, with a greater theta subsequent memory effect in controls vs PIP within anatomically defined mPFC and bilateral hippocampal ROIs. This is consistent with previous studies which have demonstrated theta subsequent memory effects in both healthy participants and people with epilepsy (Klimesch *et al*., 1996; Fell *et al*., 2011; Joensen *et al*., 2023; see Herweg *et al*., 2020 for a review). Similarly, we found greater temporal lobe theta power during the encoding of image pairs (A-B, B-C) from events where the subsequent inference (A-C) was accurately made in our control cohort, but not in the PIP group. This effect appears to originate from the temporal lobes, with a greater theta subsequent inference effect in controls vs PIP within the same anatomically defined bilateral hippocampal ROI. Again, this is in line with previous studies which have demonstrated increased theta power in the medial temporal lobe during the encoding of associations that support subsequent inference (Backus *et al*., 2016), and animal lesion studies which demonstrate that inference depends on hippocampal integrity (DeVito *et al*., 2009; van der Jeugd *et al*., 2009). Importantly, recent studies suggest that high density EEG recordings can detect oscillatory activity in deep sources such as the hippocampus (Seeber *et al*., 2019).

Consistent with our behavioural results, we found no evidence for differences in theta dynamics during the cue period immediately preceding recognition judgements between groups. In both groups, theta power averaged across all sensors was greater during the presentation of ‘old’ vs ‘new’ images, prior to their accurate recognition (i.e. for hits vs correct rejections), consistent with previous studies (Osipova *et al*., 2006). Similarly, we found little evidence for a significant difference in theta power between groups during associative memory retrieval – although we did identify a posterior central cluster of sensors that showed greater theta power during accurate memory retrieval in controls vs PIP during the first 2 s of the time window when participants were presented with four alternative forced choice options. Finally, we found that control participants – but not those with PIP – showed greater temporal lobe theta power during the accurate retrieval of inferred vs direct pairs.

The notion that reductions in theta activity seen in PIP may relate to hippocampal pathology is supported by reduced posterior hippocampal volumes described in both PIP and schizophrenia (Allebone *et al*., 2019). If PIP is indeed related to functional and/or structural abnormalities in the mTL/hippocampus, this may also explain why prevalence of psychosis is highest in temporal lobe epilepsy (Clancy *et al*., 2014). Perfusion studies during postictal psychotic episodes have also demonstrated increased perfusion in the temporal and frontal lobes (Nishida *et al*., 2006). Taken together, our findings suggest a common neural basis for integrative memory dysfunction in schizophrenia and PIP. In the context of the genomic similarities seen between participants with PIP and schizophrenia (Braatz *et al*., 2021), this study lends support to a shared mechanism predisposing to psychosis in both patient groups, although with the difference that a higher threshold of brain insult (e.g. a seizure cluster) is required to trigger psychosis in patients with PIP. The exact nature of this putative mechanism requires further elucidation, especially as this study was conducted with participants not currently experiencing an episode of psychosis.

## Acknowledgements

The authors wish to thank Rick Adams and Ingrid Martin for comments on earlier drafts of the manuscript. D. B. is supported by a UKRI Frontier Research grant (EP/X023060/1). U.V. is supported by a Wellcome Career Development Award (07306/Z/23/Z). This work is supported by researchers at the National Institute for Health and Care Research University College London Hospitals Biomedical Research Centre and Epilepsy Society. We also thank Ms Louise Price and Professor Sanjay Sisodiya for their expertise in performing TMS and use of the Epigenx database.

